# MExConn: A Mechanistically Interpretable Multi-Expert Framework for Multi-Organelle Segmentation in Connectomics

**DOI:** 10.1101/2025.11.19.689394

**Authors:** Abrar Rahman Abir, Anik Saha, Shashata Sawmya, Thomas L. Athey, Nir N. Shavit, Md Shamsuzzoha Bayzid

**Author notes:** These authors contributed equally (Abrar Rahman Abir, Anik Saha).

## Abstract

Electron microscopy (EM) provides subcellular resolution which has made it a critical tool in fields such as cellular biology and connectomics. However, manual annotation of subcellular organelles in these EM images is extremely labor-intensive and impractical at scale. While computational segmentation methods have been developed, most existing approaches are limited to segmenting a single organelle at a time, neglecting the inherent shared information present in EM images containing multiple organelles. To address this, we present MExConn, the first known interpretable multi-expert U-Net architecture in the connectomics field that employs a shared encoder and multiple decoder heads to simultaneously segment multiple organelles from the same input EM image. MExConn significantly outperforms five baselines, including single-organelle model and four state-of-the-art connectomics segmentation models in all evaluation metrics, reducing the Variation of Information by up to 33.54% on average across organelles. A key novelty of our approach is that MExConn offers mechanistic interpretability by revealing that the shared encoder learns shared representations essential for accurately segmenting multiple organelles. Through systematic analysis of encoder gradients with respect to each decoder output, we identify channel-wise importance profiles and reveal that many encoder channels are jointly essential for all organelles, while others are organelle-specific. Rigorous experiments on three connectomics datasets demonstrate the effectiveness of MExConn in both segmentation performance and interpretability, establishing it as a principled approach for multi-organelle analysis in connectomics. The source code is publicly available at https://github.com/abrarrahmanabir/MExConn.

## 1 Introduction

Connectomics is a field which seeks to map the neural wiring of the brain by reconstructing synaptic connections and cellular structures from high-resolution electron microscopy (EM) images. Segmentation of organelles, such as mitochondria, synapses, and cell membranes, is a critical step in this workflow, as it enables the identification of these structures for subsequent analysis [1]. However, achieving accurate segmentation is challenging due to the complexity of EM images, which capture nanoscale details across vast datasets. Manual annotation is prohibitively time-consuming, prone to human errors, and requires extensive proofreading, which severely limits scalability and practical feasibility [2]. Furthermore, achieving high accuracy remains difficult due to the intricate and overlapping features of organelles, necessitating advanced computational approaches.

Current automated segmentation approaches in connectomics primarily rely on architectures like 2D-UNet [3, 4, 5], 3D-UNet [6, 7], and Flood-Filling Networks (FFN) [8]. While these methods have shown promise, they often necessitate separate training for each organelle, which is computationally inefficient and limits the amount of contextual information available for learning. Training individual models for each organelle fails to exploit potential shared features across these structures, as each model learns them in isolation. In contrast, recent advances in foundation models for EM image analysis suggest that pretrained backbones can learn shared representations across diverse data, improving generalization [9, 10]. Additionally, studies in multi-task learning indicate that joint training on related tasks can enhance performance by leveraging shared features [11, 12]. These findings motivate exploring joint training strategies for organelle segmentation to improve efficiency and accuracy.

Motivated by these insights, we hypothesize that, since EM images contain multiple organelles within the same visual context, a shared encoder can learn latent representations that capture common features across these structures. These shared representations could be leveraged to segment multiple organelles simultaneously, reducing the need for separate models. Moreover, joint training on multiple organelles increases the information available to the model, as it learns from diverse labels within the same image. This approach is expected to yield better segmentation performance compared to single-organelle training, as the model can leverage correlations between organelles to refine its feature extraction and segmentation processes. Grounded in these hypotheses and observations, MExConn advances the field through two central novelties:

1. **Unified Multi-Organelle Segmentation in Connectomics via Interpretable Multi-Head U-Net:** We introduce MExConn (**M**ulti-**Ex**pert Segmentation in **Conn**ectomics), the first known interpretable multi-head U-Net architecture for connectomics, which employs a shared encoder with multiple decoder heads to simultaneously segment different organelles (mitochondria, membrane, synapses, vesicles, fusiform-vesicles, lysosomes) from the same EM image. Through rigorous experiments on three benchmark datasets, we demonstrate that our approach significantly outperforms traditional single-organelle segmentation models and state-of-the-art connectomics segmentation models.
2. **Mechanistic Interpretability of Shared Encoder Representations:** A key novelty of MExConn is that we apply mechanistic interpretability (MI) to analyze the encoder’s internal representations in our model to explore whether joint training indeed leads to shared and meaningful representations. We introduce a novel gradient-based approach for quantifying channel importance in the encoder for mechanistic interpretability in multi-head segmentation models. Specifically, for each decoder head (corresponding to each organelle), we compute the gradient of the output segmentation score with respect to each channel of the deepest encoder feature map. By averaging the absolute gradients across spatial dimensions, we obtain a quantitative measure of each channel’s importance for the segmentation of each organelle. Through this detailed analysis, comparing the most important channels across all decoder heads, we, for the first time, clearly demonstrate that the encoder learns meaningful shared representations. We show that some channels are consistently important for segmenting all organelles (shared channels), while others are uniquely important for specific organelles (organelle-specific channels), thus providing mechanistic evidence and detailed insights into how joint training enhances segmentation performance by leveraging both shared and specialized features within the encoder.

## 2 Related Work

Accurate segmentation of cellular structures in electron microscopy images is critical for connectomics. Classical approaches based on 2D and 3D U-Nets and their extensions have progressively improved segmentation performance and efficiency using dense skip connections or deeper, summation-based architectures [4, 3, 5, 7]. Affinity-based methods [13, 14] learn voxel affinities or shape descriptors and use graph agglomeration, while frameworks like GASP [6] integrate CNN outputs with hierarchical clustering. Recent work also explores multi-organellar segmentation with structural priors [15], illustrating the diversity of segmentation strategies for various organelles and spatial scales.

To reduce annotation cost and address multiple targets, recent research emphasizes unified models leveraging multi-task learning. Foundation models such as EM-DINO [9] and RETINA [10] show that large-scale pretraining on diverse EM datasets produces transferable features for multi-organelle segmentation. EM-DINO’s OmniEM decoder, for example, enables strong multi-task performance with a shared backbone. Likewise, multi-task models like Cerberus [11], MIS-Net [16], and Dense R-CNN [17] use shared encoders and attention mechanisms to segment multiple anatomical structures efficiently. Collectively, these efforts highlight the promise of multi-task learning for accurate segmentation across diverse EM targets.

While joint learning promises performance gains, understanding how such improvements arise within deep networks remains a challenge. MI methods aim to address this by analyzing and attributing model behavior to specific internal components. Network Dissection [18], for example, quantifies neuron interpretability by aligning activations with semantic concepts, enabling comparisons across models and training paradigms. More recent works aim to enhance interpretability directly within the model, as seen in SAUNet [19], which separates shape and texture processing pathways for improved robustness and transparency in medical image segmentation. In multi-task settings, [20] explored universal representations by distilling knowledge from task-specific networks into a single model, achieving state-of-the-art performance in dense prediction tasks across diverse domains. These studies highlight the growing focus on understanding feature representations obtained from the models. However, currently no existing methods applied MI to connectomics, where the complexity of organelle interactions demands interpretable models to explore shared and organelle-specific representations. Our work bridges this gap by employing MI to analyze encoder representation in joint organelle segmentation, offering novel insights into their contributions to segmentation performance.

## 3 Methodology

### 3.1 Problem Formulation

Let *X* ⊆ ℝ_*H×W*_ denote the space of electron microscopy images, where each image *x* ∈ *X* is a grayscale image of size *H* × *W*. Let *Y* ⊆ {0, 1}^*K×H×W*^ represent the space of binary segmentation masks, where *K* denotes the number of distinct organelles of interest. Each segmentation mask *y* ∈ *Y* consists of *K* binary channels, *y* = (*y*^(1)^, *y*^(2)^, …, *y*^(*K*)^), where each channel *y*^(*k*)^ ∈ {0, 1}^*H×W*^ corresponds specifically to the binary mask for the *k*-th organelle. We are provided with a dataset 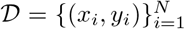, where *x*_*i*_ ∈ *X* is an EM image, and the corresponding segmentation masks are defined as: 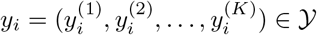.

The objective is to learn a function: *F*: *X* → *Y*, such that *F* accurately predicts the segmentation masks, identifying pixels associated with each of the *K* organelles in the EM images.

### 3.2 MExConn Joint Training Architecture

The MExConn architecture is a UNet-inspired model which comprises a single encoder and *K* decoder heads, with the encoder generating shared feature representations that are passed to each decoder to produce specialized segmentation outputs.

The encoder is a function: 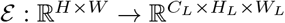, where *C*_*L*_ is the number of channels in the final feature map, and *H*_*L*_, *W*_*L*_ are the spatial dimensions after *L* downsampling operations. The encoder consists of *L* stacked convolutional blocks, each followed by a max-pooling operation to reduce spatial dimensions. Each convolutional block consists of two convolutional layers with batch normalization and ReLU activations, followed by a Squeeze-and-Excitation (SE) module. A residual connection adds the block’s input to its output. Dropout is applied after the first convolution to promote regularization. Each block transforms an input feature map to produce an output feature map 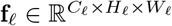, where *C*_**l**_ is the number of channels, and *H*_**l**_, *W*_**l**_ are the spatial dimensions after **l** max-pooling operations. From the encoder, we thus extract a set of hierarchical feature maps {**f**_1_, **f**_2_, …, **f**_*L*_},.

Each of the *K* decoder heads, denoted *D*_*k*_, maps the set of encoder features{**f**_1_, …, **f**_*L*_*}* to a segmentation mask *y*^(*k*)^ ∈{0, 1} ^*H×W*^ for the *k*-th organelle. Each decoder employs a UNet-style upsampling path with *L* −1 stages, where each stage upsamples the feature map and combines it with the corresponding encoder feature map **f**_**l**_ via skip connections, followed by convolutional processing to refine features. The final stage applies a 1× 1 convolution and sigmoid activation to produce the binary segmentation mask *y*^(*k*)^. Figure 1 presents the joint training architecture of MExConn.

**Figure 1:**
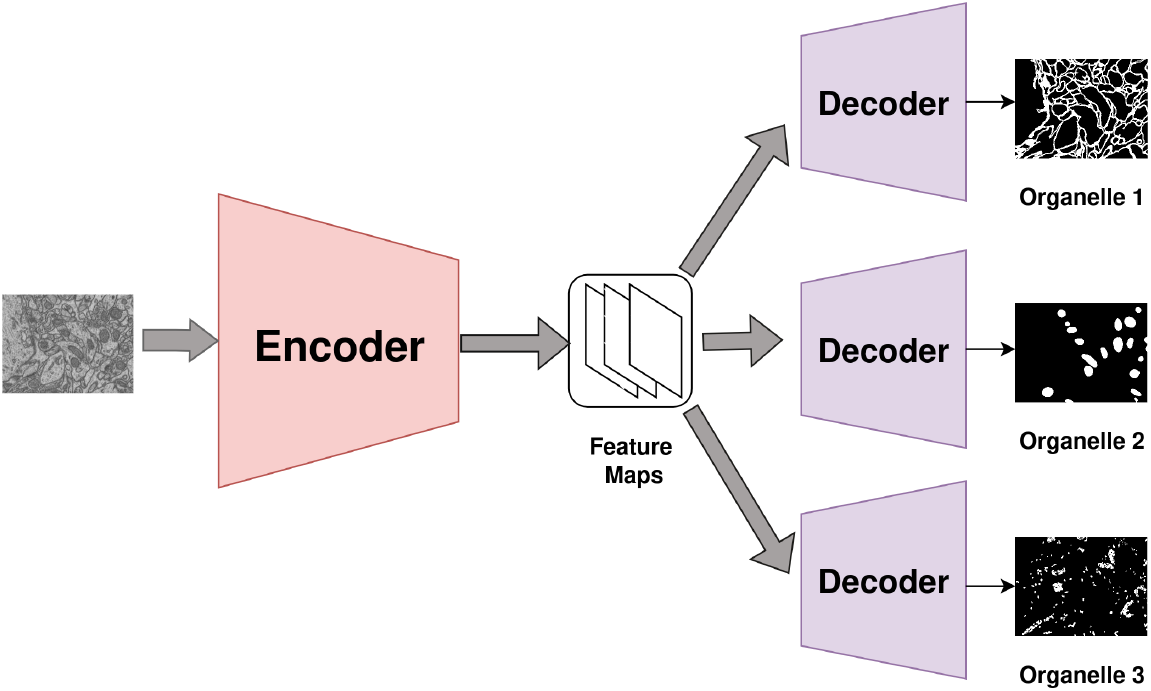
Overview of the MExConn architecture: A shared encoder extracts feature maps from an input EM image. These feature maps are then passed to multiple decoder heads, each specialized for segmenting a specific organelle, enabling simultaneous multi-organelle segmentation from a single input image.

### 3.3 Training Objective

The training objective is to optimize the segmentation performance across all *K* organelles for a given input electron microscopy (EM) image *x*_*i*_ ∈ *X* and its corresponding ground truth masks 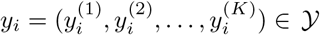. The predicted masks are given by 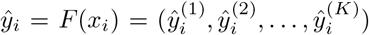, where *F*: *X* → *Y* is the segmentation function. We use a composite loss function combining Dice loss, Focal loss, and Binary Cross-Entropy (BCE) loss, defined as: ℒ (ŷ_*i*_, *y*_*i*_) = ℒ_Dice_(ŷ_*i*_, *y*_*i*_) + ℒ_Focal_(ŷ_*i*_, *y*_*i*_) + ℒ_BCE_(ŷ_*i*_, *y*_*i*_)

### 3.4 Computing Channel Importance Through Mechanistic Interpretability

To identify whether the encoder learns shared representations which is necessary for multi-organelle segmentation, we want to investigate the sensitivity of decoder outputs or logits with respect to each channel of the encoder representation. Therefore, we compute channel importance scores for each decoder head, providing a mechanistic interpretability framework to analyze the learned representations.

The process begins with an input image patch passed through the U-Net encoder, yielding a deepest feature map *f*_*L*_ of shape (*C*_*L*_, *H*_*L*_, *W*_*L*_), where *C*_*L*_ denotes the number of channels, and *H*_*L*_ and *W*_*L*_ represent height and width of the feature map *f*_*L*_, respectively. This feature map is fed into multiple organelle-specific decoder heads. For each decoder head *D*_*k*_ corresponding to organelle *k*, the output logits are computed. We aggregate these logits spatially by summing over all pixels, resulting in a scalar score *S*_*k*_ = ∑_*h,w*_ logits_*k*_(*h, w*), which quantifies the overall logit score for organelle *k*’s segmentation.

Next, we perform backpropagation to compute the gradient of *S*_*k*_ with respect to *f*_*L*_, denoted as 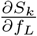, a tensor of shape (*C*_*L*_, *H*_*L*_, *W*_*L*_). This gradient reflects the sensitivity of the output score to each channel at every spatial location. To derive a per-channel importance measure, we take the absolute value of this gradient, 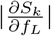 and average it over the spatial dimensions *H*_*L*_ and *W*_*L*_, yielding a vector *I*_*k*_ of length *C*_*L*_, where each element

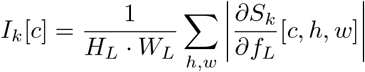

represents the importance of channel *c* for organelle *k*. Figure 2 summarizes the approach of computing channel importance via saliency analysis.

**Figure 2:**
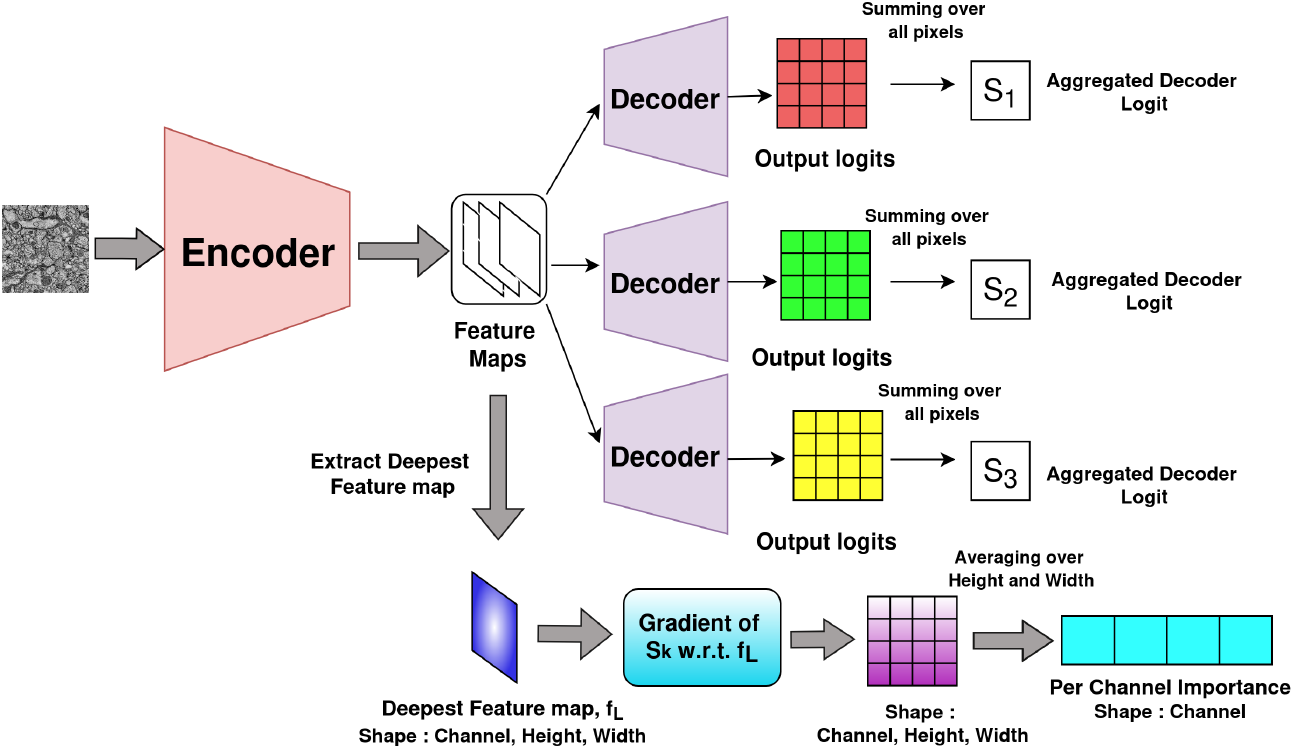
Mechanistic interpretability analysis of the MExConn architecture. After extracting feature maps from the input EM image using a shared encoder, the deepest encoder feature map (*f*_*L*_) is isolated. For each decoder head, output logits are summed across all pixels to yield an aggregated decoder logit (*S*_*k*_). The gradient of *S*_*k*_ with respect to *f*_*L*_ is computed and averaged over spatial dimensions to obtain per-channel importance values.

## 4 Experimental Results

We performed an extensive experimental analysis to validate MExConn on three benchmark datasets. We first outline the experimental setup and datasets, followed by results, showing that joint training across organelles in MExConn consistently outperforms single-organelle training. We further analyze MExConn through mechanistic interpretability, showing that the encoder captures both shared and organelle-specific features, which are essential for accurate segmentation. Additionally, we investigate the effectiveness in 3D volumetric segmentation and present a case study on the most challenging test samples to demonstrate the efficacy of our approach qualitatively.

### 4.1 Experimental Setup

We evaluated MExConn across three distinct EM datasets. Two of them are connectomics EM datasets: Multi Class Annotated Mouse CA1 Hippocampus (Multiclass-EPFL) [21], and Drosophila Ventral Nerve Cord (Drosophila-VNC) [22]. We also use Mouse Urinary Bladder dataset (Urocell) [23] which is widely used in EM organelle segmentation and reconstruction analysis [10, 24, 25]. Each of the datasets contains segmentation masks across three different organelles for each EM image. The following Datasets section presents further details of the datasets. We refer to MExConn training strategy as joint training because a single model jointly segments multiple organelles from the same input image by leveraging shared representations across organelles.

#### Baselines

To extensively evaluate the segmentation performance, we compare MExConn against five representative baselines, organized into three categories:

1. **Single-organelle Models:** We consider single-organelle training as a baseline. In single-organelle training, the model is trained to segment only a particular organelle, unlike MExConn.
2. **Connectomics-specific segmentation models:** *PyTorch Connectomics (PyTC)* [26] is a widely-used framework for EM segmentation in connectomics. *FusionNet* [27] is a connectomics-specific convolutional model for EM segmentation. *RETINA* [28] is another baseline pretrained on largescale EM images and fine-tuned for task-specific segmentation.
3. **Foundation models:** *Segment Anything Model (SAM)* [29] is a promptable, general-purpose foundation segmentation model.

#### Implementation Details

MExConn is trained for 20 epochs with a batch size of 8. We focus on the task of segmenting multiple organelles from two-dimensional EM image, an essential step in the connectomics pipeline. We use Variation of Information (VI), Dice coefficient, Intersection over Union (IoU), Precision, and Recall as our evaluation metrics. All experiments are conducted five times and on a cluster equipped with four NVIDIA RTX A4500 GPUs. Hyperparameters for MExConn were optimized using Ray Tune [30] with an ASHA scheduler [31]. All tuning decisions were based solely on the validation set, and the final hyperparameter configuration was fixed before evaluating on the test set. We report all final hyperparameters in the Supplementary Material. To ensure fairness, all baseline models were run on the same hardware and with identical data splits and we used the default hyperparameters exactly as provided by the authors.

### 4.2 Datasets

The three datasets utilized in this study are summarized in Table 1. For each dataset, we apply a 70:15:15 split to create training, validation, and test sets, respectively. In all experiments, we adopt a patch-based training approach. Each EM image is partitioned into overlapping patches of 256 ×256 pixels, with a stride of 128 pixels between adjacent patches. Images and their corresponding segmentation masks are converted to grayscale during preprocessing. Additionally, mask patches are binarized to maintain consistent foreground-background labeling.

**Table 1:** Summary of organelle segmentation datasets used in this study. The **Train** and **Test** columns indicate the number of image patches extracted from each dataset split.

| Domain | Species | Organelle | Train | Test |
| --- | --- | --- | --- | --- |
| Urocell [23] | Mouse | fusiform-vesicles | 716 | 155 |
|  |  | lysosomes | 716 | 155 |
|  |  | mitochondria | 716 | 155 |
| Multiclass-EPFL [21] | Mouse | membranes | 1645 | 385 |
|  |  | mitochondria | 1645 | 385 |
|  |  | vesicles | 1645 | 385 |
| Drosophila-VNC [22] | *Drosophila* | membranes | 686 | 147 |
|  |  | mitochondria | 686 | 147 |
|  |  | synapses | 686 | 147 |

### 4.3 MExConn Yields Superior Segmentation Performance Across All Baseline Methods

To rigorously assess the benefit of joint training, we compare segmentation performance between models trained using single-organelle strategy, our proposed joint training approach MExConn, and four other baselines PyTC, FusionNet, RETINA and SAM. We aim to empirically validate whether joint training leads to superior segmentation accuracy. For the single training approach, we train independent models for each organelle using only its corresponding annotations. In contrast, our joint training employs a shared encoder with multiple organelle-specific decoders, allowing for the simultaneous segmentation of multiple organelles. However, the encoder and decoder components in both the approaches share similar architectures. PyTC and FusionNet are trained from scratch on each organelle. RETINA is finetuned on our training splits, and inference is performed using these finetuned models. Since SAM is a promptable foundation model, we provide it with a small subset of positive and negative pixel masks as prompts. All the baselines are trained or finetuned with single organelle masks corresponding to images in contrast to our joint training approach. Afterwards we evaluate all models on the respective test splits. The results, summarized in Table 2,3, and 4, reveal that MExConn consistently achieves the highest Dice, IoU, and Precision scores, indicating more accurate segmentations across all domain-organelle pairs. MExConn also attains lower VI scores than single-organelle training and most other baselines, with the only exception being synapse segmentation in Drosophila-VNC. SAM attains the highest Recall in many cases but at the cost of substantially lower Precision, reflecting its tendency to over-segment by predicting a large proportion of pixels as positive. In terms of VI, joint training outperforms single organelle training by a margin ranging from 9.34% (vesicles in Multiclass) to 59.93% (lysosomes in Urocell) across all domain-organelles combinations, only except for synapses in Drosophila-VNC where joint training yields a higher VI of 0.1012 compared to 0.0481 in case of single organelle training. Unlike other organelles, synapses are comparatively harder to segment [32] because they are rare, extremely sparse and show high structural variability in EM images [33]. Hence, synapses are likely to be better captured by single-organelle training, which can more effectively focus on their unique patterns. Averaged across all domains and organelles, MExConn yields a 10.4% reduction in VI compared to single-organelle models. Assessing domain-specific gains, we observe that in Urocell-3, the largest improvement is for lysosomes and the smallest for fusiform vesicles. In Multiclass-EPFL, mitochondria benefit the most and vesicles the least. For Drosophila-VNC, mitochondria again see the highest gain, while synapses show a decrease with joint training. The highest domain-wise average gain is observed in Urocell, where joint training reduces VI by 33.54% on average across organelles, highlighting the benefits of leveraging shared contextual information in the segmentation task.

**Table 2:** Segmentation performance for Urocell. Best values per metric and organelle are in bold.

| Method | Fusiform-Vesicles |  |  |  |  | Lysosomes |  |  |  |  | Mitochondria |  |  |  |  |
| --- | --- | --- | --- | --- | --- | --- | --- | --- | --- | --- | --- | --- | --- | --- | --- |
|  | Dice | VI | IoU | Precision | Recall | Dice | VI | IoU | Precision | Recall | Dice | VI | IoU | Precision | Recall |
| Single | 0.554 | 0.879 | 0.404 | 0.553 | 0.568 | 0.315 | 0.208 | 0.234 | 0.306 | 0.693 | 0.785 | 0.291 | 0.675 | 0.747 | 0.847 |
| MExConn | **0.754** | 0.782 | **0.611** | **0.67** | 0.868 | **0.835** | **0.083** | **0.789** | **0.948** | **0.817** | **0.94** | 0.205 | **0.897** | **0.942** | 0.939 |
| PyTC | 0.4927 | **0.5073** | 0.343 | 0.3713 | 0.8192 | 0.3478 | 0.6522 | 0.2658 | 0.3778 | 0.3833 | 0.7923 | 0.2077 | 0.6724 | 0.7985 | 0.816 |
| FusionNet | 0.3455 | 0.8807 | 0.216 | 0.2118 | 0.9378 | 0.1471 | 0.2214 | 0.0936 | 0.0838 | 0.6011 | 0.0106 | **0.1912** | 0.0054 | 0.1395 | 0.0055 |
| RETINA | 0.3045 | 0.9852 | 0.1904 | 0.1866 | 0.8265 | 0.1254 | 0.254 | 0.0798 | 0.0714 | 0.5124 | 0.0092 | 0.2173 | 0.0047 | 0.1205 | 0.0048 |
| SAM | 0.3445 | 1.1859 | 0.2186 | 0.2208 | **0.9742** | 0.0918 | 0.8925 | 0.052 | 0.0542 | 0.6124 | 0.2901 | 0.9126 | 0.1764 | 0.1779 | **0.9719** |

**Table 3:** Segmentation performance for Multiclass. Best values per metric and organelle are in bold.

| Method | Membranes |  |  |  |  | Mitochondria |  |  |  |  | Vesicles |  |  |  |  |
| --- | --- | --- | --- | --- | --- | --- | --- | --- | --- | --- | --- | --- | --- | --- | --- |
|  | Dice | VI | IoU | Precision | Recall | Dice | VI | IoU | Precision | Recall | Dice | VI | IoU | Precision | Recall |
| Single | 0.799 | 0.952 | 0.682 | 0.741 | 0.904 | 0.67 | 0.438 | 0.577 | 0.635 | 0.834 | 0.679 | 0.323 | 0.525 | 0.628 | 0.788 |
| MExConn | **0.852** | 0.829 | **0.759** | **0.809** | 0.93 | **0.905** | 0.311 | **0.853** | **0.9** | 0.941 | **0.742** | 0.293 | **0.603** | **0.689** | 0.848 |
| PyTC | 0.5807 | **0.4193** | 0.4286 | 0.4603 | 0.8919 | 0.2099 | 0.7901 | 0.1202 | 0.1202 | **0.9968** | 0.3595 | 0.6405 | 0.2239 | 0.2294 | 0.9282 |
| FusionNet | 0.7505 | 0.6087 | 0.6017 | 0.7429 | 0.7583 | 0.7498 | **0.2759** | 0.6018 | 0.735 | 0.7652 | 0.0056 | **0.1705** | 0.0028 | 0.6407 | 0.0028 |
| RETINA | 0.6614 | 0.6809 | 0.5302 | 0.6547 | 0.6683 | 0.6392 | 0.3166 | 0.513 | 0.6266 | 0.6523 | 0.0049 | 0.1938 | 0.0024 | 0.5531 | 0.0024 |
| SAM | 0.3641 | 1.1766 | 0.2235 | 0.2251 | **0.9708** | 0.2196 | 1.2431 | 0.1242 | 0.1244 | 0.9835 | 0.1025 | 1.0192 | 0.0541 | 0.0541 | **0.9947** |

**Table 4:** Segmentation performance for Drosophila-VNC. Best values per metric and organelle are in bold.

| Method | Membranes |  |  |  |  | Mitochondria |  |  |  |  | Synapses |  |  |  |  |
| --- | --- | --- | --- | --- | --- | --- | --- | --- | --- | --- | --- | --- | --- | --- | --- |
|  | Dice | VI | IoU | Precision | Recall | Dice | VI | IoU | Precision | Recall | Dice | VI | IoU | Precision | Recall |
| Single | 0.759 | 1.059 | 0.617 | 0.661 | 0.914 | 0.429 | 0.554 | 0.319 | 0.414 | 0.714 | 0.484 | 0.048 | 0.475 | 0.476 | 0.475 |
| MExConn | **0.876** | 0.789 | **0.781** | **0.867** | 0.888 | **0.77** | 0.408 | **0.694** | **0.875** | 0.789 | **0.649** | 0.101 | **0.593** | **0.902** | 0.628 |
| PyTC | 0.6804 | **0.3196** | 0.5167 | 0.8433 | 0.5719 | 0.6116 | 0.3884 | 0.4459 | 0.7213 | 0.5331 | 0.4553 | 0.4588 | 0.4289 | 0.714 | 0.3341 |
| FusionNet | 0.7066 | 0.7174 | 0.5466 | 0.5762 | 0.9133 | 0.5028 | **0.2851** | 0.3359 | 0.6276 | 0.4194 | 0.0718 | **0.039** | 0.0373 | 0.3735 | 0.0397 |
| RETINA | 0.6227 | 0.8026 | 0.4817 | 0.5078 | 0.8048 | 0.4286 | 0.3271 | 0.2864 | 0.535 | 0.3575 | 0.062 | 0.0443 | 0.0322 | 0.3225 | 0.0343 |
| SAM | 0.3578 | 1.1709 | 0.2179 | 0.219 | **0.979** | 0.2242 | 1.1859 | 0.1268 | 0.1284 | **0.9259** | 0.0226 | 1.0258 | 0.0115 | 0.0115 | **0.991** |

#### Statistical Significance

All improvements of our method over all the baselines are statistically significant, as determined by a paired two-sided t-test (*p <* 0.05) across five independent runs. This confirms that the observed performance gains are unlikely to be due to random variation and reflect a consistent advantage of our approach across experimental runs.

### 4.4 Mechanistic Interpretability in MExConn Reveals Encoder Learns Both Shared and Organelle-Specific Representations

We aim to investigate whether the encoder in MExConn, when jointly trained for multiclass organelle segmentation, learns shared and organelle-specific representations. To this end, we analyze the final encoder activations using mechanistic interpretability. Specifically, for each organelle head, we compute channel importances following the procedure outlined in the Methodology section. We then rank the encoder channels in descending order of their corresponding importance scores and select the top 100 channels per organelle. By comparing these sets, we quantify how many channels are shared versus uniquely specialized. As shown in Figure 3, some encoder channels emerge consistently across all three organelles, indicating shared representations, while others appear exclusively for individual organelles, suggesting specialization. Among the three datasets, the Multiclass-EPFL domain exhibits the highest number of shared channels (16) among the top 100 channels per organelle, while Urocell shows the lowest (8). Correspondingly, Multiclass-EPFL has the lowest count of total organelle-specific channels (112), reflecting more integrated representations. In contrast, Urocell has the highest total number of organelle-specific channels (134), implying more task-specific encoding. At a finer granularity, mitochondria segmentation in Urocell relies most heavily on unique channels (48), whereas the membrane-specific representation in Multiclass-EPFL uses the fewest (31). These findings validate that joint training induces a mix of shared and specialized features within the encoder. In the following subsection, we evaluate the relative importance and contribution of the shared and organelle-specific channels to segmentation accuracy.

**Figure 3:**
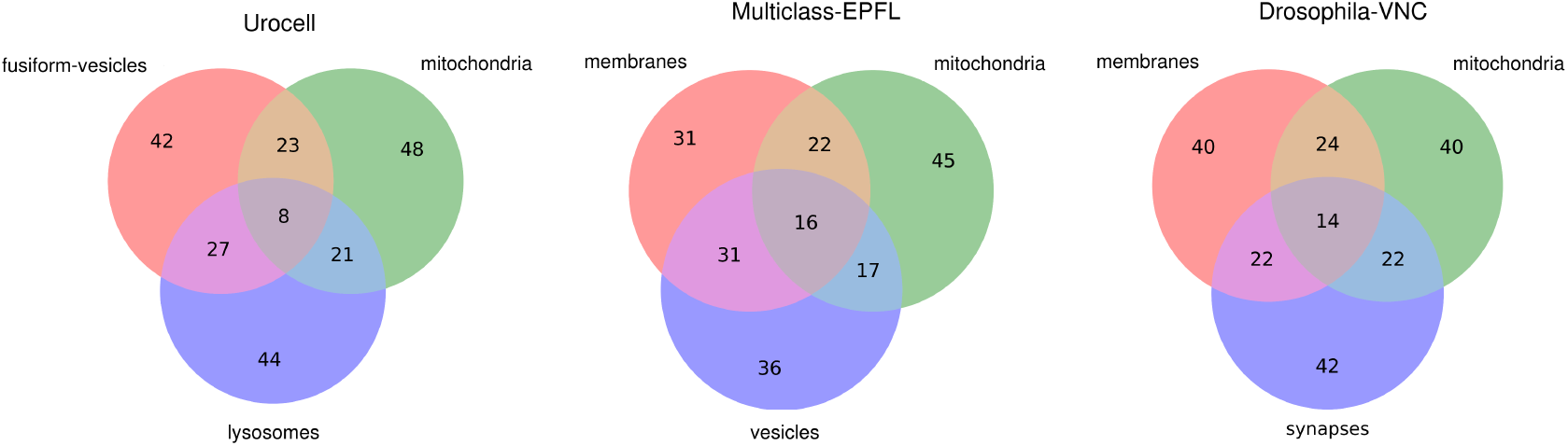
Analysis of the 100 most salient encoder channels for each organelle across all datasets. The overlap counts in these Venn diagrams indicate the number of shared (across organelles) and organelle-specific channels.

### 4.5 Encoder Learns Shared Representations Essential for Accurate Multi-Organelle Segmentation

We want to investigate whether the learned encoder representations encode shared and organelle-specific information in a functionally meaningful way. Specifically, we seek to empirically validate the role and relative contribution of shared and organelle-specific channels to segmentation performance. To this end, we conduct this experiment where different sets of encoder channels are selectively removed and the impact is measured. We begin with the joint-trained model and ablate the activations of all shared channels by setting their output activations to zero. We then perform a similar ablation for the organelle-specific channels, where all channels unique to any decoder head are zeroed out. For comparison, we also randomly selected 16, 32, 64, and 128 channels (out of 256 total) and dropped them in the same way and repeated this experiment five times for reliability. In each case, we report the percentage increase in Variation of Information (VI) relative to the original joint-trained network, indicating degradation in performance due to information loss. As visualized in Figure 5, removal of shared channels consistently results in higher VI degradation than random removal, and the effect is even more pronounced for organelle-specific channel ablation. The joint-trained network achieves optimal segmentation performance when no channels are ablated. Progressively removing up to 128 random channels (50% of 256) has minimal impact, with little to no degradation up to 64 channels. However, ablating just 8, 14, or 16 shared channels (less than 5% of 256) in these three datasets causes a substantial drop in performance, highlighting their critical role in capturing essential features. Averaged across all datasets and organelles, ablation of shared channels leads to a 170.05% increase in VI, indicating substantial loss of segmentation fidelity. In contrast, progressive random ablation causes much smaller average performance drops of 56.71% (for 128 channels), 12.74% (for 64 channels), 3.89% (for 32 channels), and 0.82% (for 16 channels), respectively. The effect is even more severe for organelle-specific channels, with an average VI increase of 212.37%. This trend holds across all datasets and organelles except for the segmentation of synapses in Drosophila-VNC. Notably, synapse segmentation in Drosophila-VNC is the only case where ablating shared or organelle-specific channels results in a smaller degradation than removing 128 random channels. Mechanistically, this suggests that for synapses, which are sparse and highly variable in EM images as noted by [33], the model does not rely on a small, functionally critical set of encoder channels. Instead, synapse-relevant information appears to be distributed more diffusely across many channels. As a result, ablating a large number of random channels can disrupt the already limited synapse-representing capacity more than removing those channels identified as shared or organelle-specific. Consistent with this mechanistic insight, as shown in Table **??**, synapse segmentation is the only case where joint training did not outperform single-organelle training. This further highlights the value of our mechanistic interpretability, which reveals how critical features are distributed for the organelles. Next, we analyze the number of channels ablated in each case and their corresponding relative impact on segmentation accuracy across datasets. In the Urocell-3 dataset, the ablation of just 8 shared channels increases VI by 225.64%, compared to only 48.23% when randomly ablating as many as 128 out of 256 channels. Organelle-specific ablation (removing 134 channels) in the same setting causes a 277.64% VI increase.

Similarly, for Multiclass-EPFL, removing the 112 organelle-specific channels leads to a 176.06% degradation on average, substantially greater than the 23.04% caused by removing 128 random channels. The highest degradation is observed for mitochondria segmentation in Urocell-3, 380.22% for shared channel ablation and 517.1% for organelle-specific. Even the lowest drops across all ablations remain non-trivial, 78.9% in synapse segmentation for shared channels in Drosophila-VNC and 96.11% for fusiform vesicles in Urocell-3 under organelle-specific ablation. To analyze the impact of eliminating these organelle-specific channels at a finer granularity, we evaluate the performance degradation caused by progressively ablating 5, 10, 15, and so on channels specific for each organelle within each domain. Figure 4 illustrates the resulting gradually increasing trend of VI in most cases, highlighting the cumulative effect of this progressive ablation.

**Figure 4:**
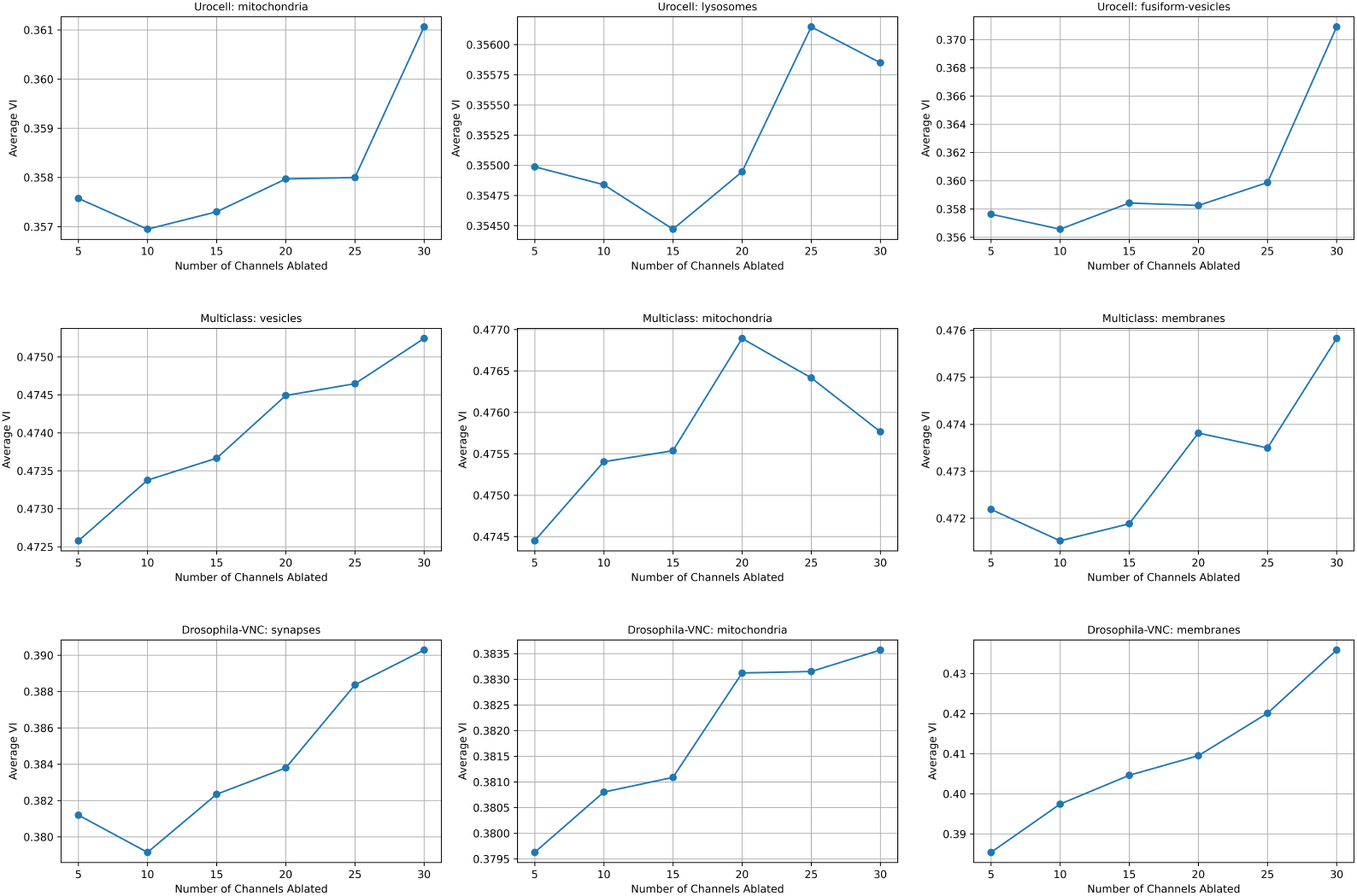
Impact of progressive removal of organelle-specific channels on segmentation performance (VI) across all datasets and organelles.

**Figure 5:**
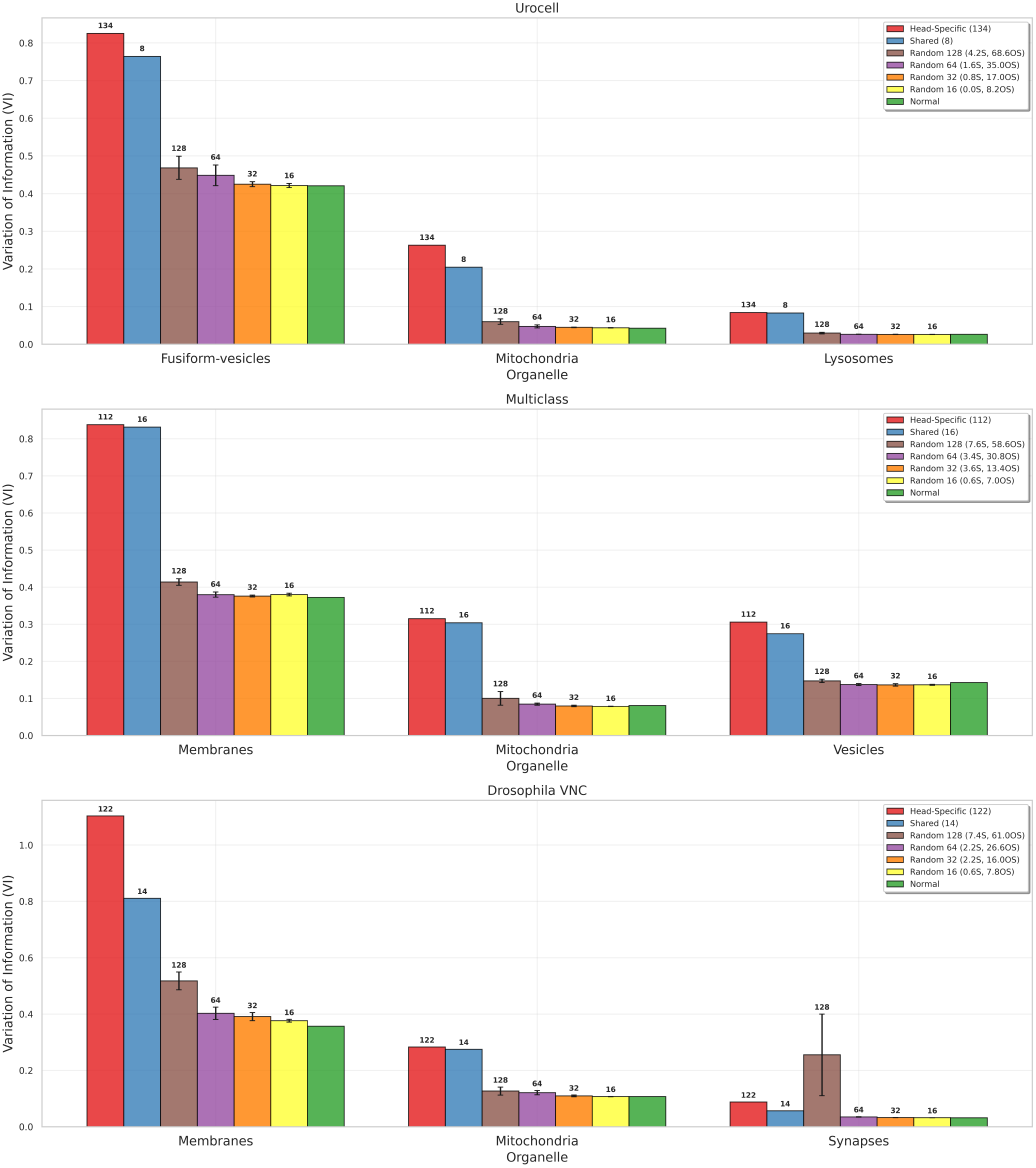
Comparison of segmentation performance (VI) on ablation of shared, head-specific, and random channels for each organelle across all datasets. The full model (no ablation; 256 channels) is included as a baseline. Bar labels above each bar indicate the number of channels removed. For random ablation, the color legends show the mean number of shared (S) and organelle-specific (OS) channels present in randomly selected channel sets (displayed in parentheses).

To further investigate the contributions of shared and organelle-specific channels, we analyze the composition of randomly ablated channels across five experimental runs, as illustrated in Figure 5 (see the average number of shared and organelle-specific channels among the random ones, shown in color legends). Specifically, we compute the mean number of shared and organelle-specific channels removed during random ablations of 16, 32, 64, and 128 channels. This analysis reveals a notable resilience in the joint-trained model. For instance, in the Urocell-3 dataset, ablating all 8 shared channels results in a substantial VI increase of 225.64%. In contrast, random ablation of 128 channels, which also includes approximately 4 shared channels on average, yields a markedly lower VI increase of 48.23%. Similarly, ablating all 134 organelle-specific channels in Urocell-3 causes a 277.64% VI increase, yet random ablation of 128 channels, containing roughly 68 organelle-specific channels on average, results in significantly less degradation (48.23%). This observation suggests a degree of informational redundancy among shared and organelle-specific channels. The encoder’s performance remains robust when a substantial portion of these channels is removed, as their complementary roles allow partial compensation for the loss. However, complete ablation of either shared or organelle-specific channels leads to a catastrophic loss of essential information, severely impairing segmentation accuracy.

### 4.6 Beyond 2D: MExConn Retains its Effectiveness in 3D Volumetric Segmentation

Having thoroughly established the effectiveness of MExConn for 2D multi-organelle segmentation, we next sought to investigate whether its advantages persist in the context of 3D volumetric data. Specifically, we aimed to assess whether the benefits of joint training, demonstrated in 2D, extend to 3D, where structural complexity and contextual cues across slices present additional challenges. This experiment was conducted on the Drosophila-VNC 3D electron microscopy volume, comprising 1024 ×1024 ×20 voxels. For training and testing, we extracted non-overlapping 64^3^ volumetric patches and performed a train-test split. As shown in Table 5, MExConn continues to outperform single training and all baselines for all organelles in the 3D setting. The results indicate that joint training not only scales to volumetric segmentation but also delivers consistent improvements over specialized, organelle-specific models. This confirms the generalizability of MExConn’s joint learning strategy for biological segmentation in both 2D and 3D domains.

**Table 5:** Variation of Information (VI) for MExConn and baselines across organelles. Lower VI indicates better segmentation.

| Method | Membranes | Mitochondria | Synapses |
| --- | --- | --- | --- |
| MExConn | **0.8980** | **0.3281** | **0.0245** |
| Single Training | 0.9076 | 0.5570 | 0.3937 |
| FusionNet | 0.9691 | 0.7682 | 0.4628 |
| RETINA | 0.9848 | 0.8150 | 0.5011 |

### 4.7 Case Study

To qualitatively evaluate the efficacy of our proposed MExConn framework, we conduct a detailed case study on the most challenging samples from each dataset. For every dataset, we identified the test image that was the most difficult to segment in terms of prediction accuracy. We then visualized and compared the segmentation results obtained from single organelle training and our joint multiorganelle segmentation MExConn. As shown in Figure 6, the single organelle models often yield porous or fragmented membrane predictions, miss significant organelle structures, and produce false positives. In contrast, MExConn produces more accurate, contiguous segmentations and effectively captures even the subtle or ambiguous organelles. These visualizations clearly demonstrate the superiority of our joint training approach, highlighting its effectiveness in the most challenging cases.

**Figure 6:**
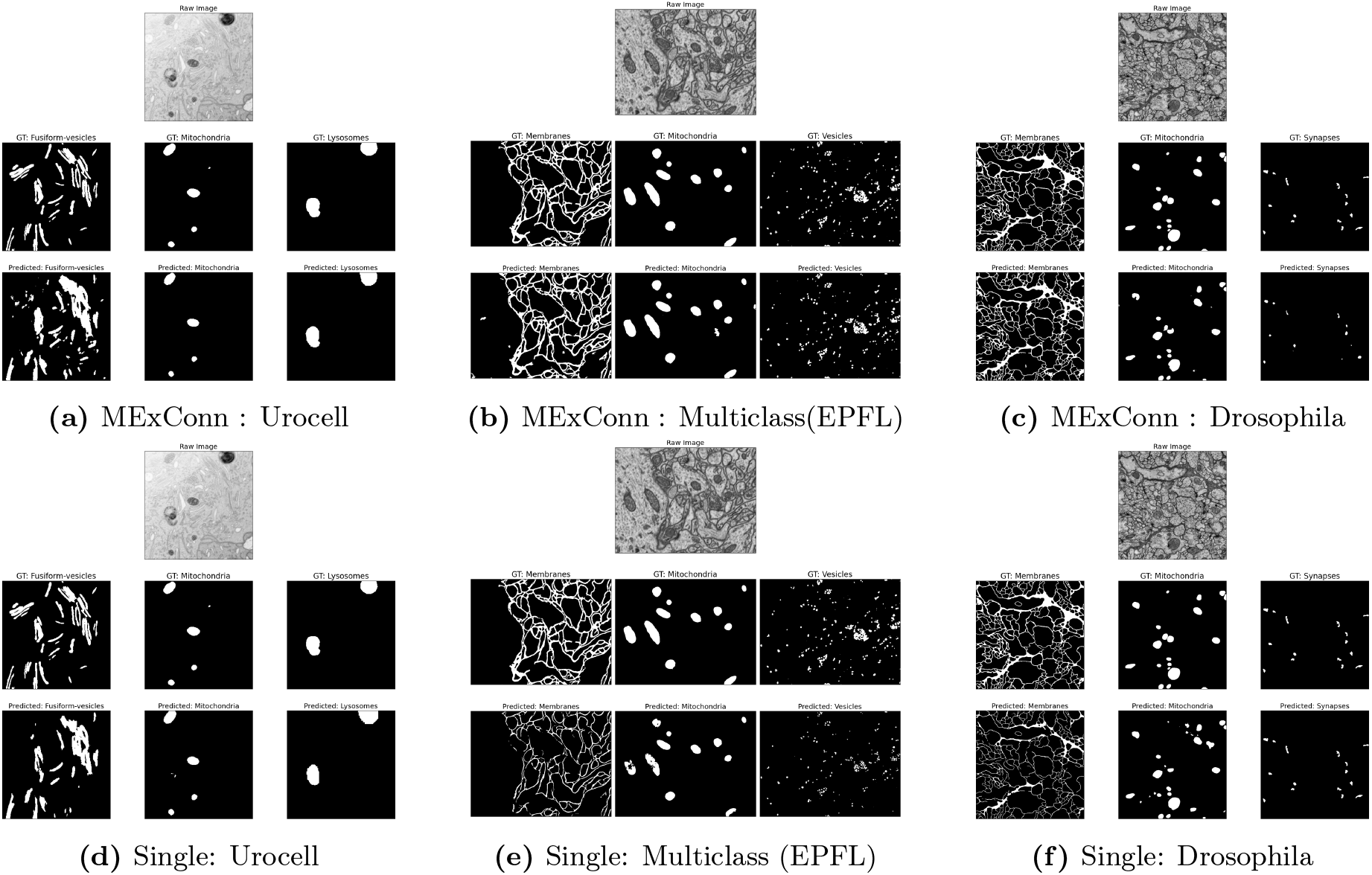
Visual comparison of segmentation outputs. Top row: MExConn (joint training) results for three datasets. Bottom row: Corresponding single-organelle segmentation results for each dataset. Joint training with MExConn achieves visibly superior segmentation for the organelles. We show the input EM image, Ground truth (GT) and predicted segmentation mask.

## 5 Conclusion

We introduced MExConn, a novel multi-expert U-Net framework for connectomics that enables simultaneous multi-organelle segmentation from EM images using a shared encoder and multiple organelle-specific decoder heads. Our approach achieves superior segmentation accuracy compared to single-organelle models across three diverse datasets, demonstrating that joint training leverages shared information to improve the performance. Remarkably, MExConn provides mechanistic interpretability through gradient-based channel importance analysis and reveals how the encoder learns both shared representations essential for all organelles and organelle-specific features critical for specialized segmentation. Furthermore, MExConn retains its effectiveness in 3D volumetric segmentation as well.

Despite these strengths, our study has several limitations. First, the current model supports a fixed, predefined set of organelle classes, which constrains its applicability to previously unseen structures. Second, the approach assumes the availability of sufficient annotation for each organelle type, limiting its use in low-annotation regimes and unlabeled domains. Therefore, extending MExConn to support a dynamic set of organelles could address the novel organelles. Additionally, integrating semi-supervised or self-supervised learning frameworks may reduce reliance on labeled data and improve scalability. We also plan to explore domain adaptation and active learning pipelines to further enhance cross-domain generalization.

Overall, MExConn lays the groundwork for an interpretable, efficient, and scalable approach to multiorganelle segmentation. As connectomics datasets continue to grow in scale and complexity, models like ours that balance accuracy, mechanistic insight, and adaptability will be essential.

## Notes

### Competing Interest Statement

The authors have declared no competing interest.

