## Supplementary Table 1 for "MExConn: A Mechanistically Interpretable Multi-Expert Framework for Multi-Organelle Segmentation in Connectomics"

### 1 Hyperparameter Details of MExConn

The hyperparameter details of MExConn are presented in Table S1.

### 2 Evaluation Metrics

To rigorously evaluate segmentation performance in electron microscopy (EM) images, we employ a set of widely-used metrics. These metrics capture both region-level overlaps and boundary-level disagreements between the predicted and ground-truth segmentations.

#### 2.1 Dice Score

The Dice Score quantifies the overlap between the predicted segmentation mask $P$ and the ground truth mask $G$:

$$\text{Dice}(P, G) = \frac{2|P \cap G|}{|P| + |G|} = \frac{2TP}{2TP + FP + FN} \tag{1}$$

where $TP$, $FP$, and $FN$ represent the number of true positives, false positives, and false negatives, respectively. A Dice score of 1 indicates perfect overlap. Since, we are evaluating on a binary segmentation task, Dice Score is also equivalent to F1-score.

#### 2.2 Precision

Precision quantifies the proportion of predicted positive pixels that are actually correct:

$$\text{Precision} = \frac{TP}{TP + FP} \tag{2}$$

High precision implies that the model produces few false positives, which is crucial when minimizing over-segmentation errors in fine structures.

Table S1: Summary of Model and Training Hyperparameters

| **Hyperparameter** | **Value/Default** | **Description** |
|---|---|---|
| *Dataset / Training* | | |
| `data_root` | data | Root directory for datasets |
| `domain` | - | Dataset subfolder/domain (e.g., drosophila-vnc, multiclass) |
| `out_path` | - | Path to save trained model checkpoints |
| `patch_size` | 256 | Size (in pixels) of image/mask patches extracted for training |
| `stride` | 128 | Stride (in pixels) between patch extractions |
| `batch_size` | 8 | Batch size for DataLoader |
| `epochs` | 20 | Number of training epochs |
| `lr` | $1 \times 10^{-4}$ | Learning rate for optimizer (Adam) |
| `device` | cuda/cpu | Computation device selection |
| `num_workers` | 4 | Number of worker processes for DataLoader |
| *Model Architecture* | | |
| `in_ch` | 1 | Number of input channels (grayscale) |
| `base_features` | [32,64,128,256] | List of feature sizes for encoder |
| `out_ch_per_head` | 1 | Number of output channels per decoder head |
| `num_heads` | 3 | Number of decoder heads (one per organelle) |
| `p_drop` | 0.2 | Dropout probability in ResidualDoubleConv and decoders |
| `use_se` | True | Use Squeeze-and-Excitation blocks (SEBlock) |
| `reduction` (SE) | 16 | Channel reduction ratio in SEBlock |
| *Loss Functions* | | |
| Dice loss smooth | 1.0 | Smoothing constant for Dice loss |
| Focal loss $\alpha$ | 0.8 | Focal loss weighting parameter |
| Focal loss $\gamma$ | 2 | Focal loss focusing parameter |
| BCEWithLogitsLoss | – | Used as-is (no custom hyperparams) |

### 2.3 Recall

Recall measures the fraction of actual positive pixels that are correctly predicted:

$$\text{Recall} = \frac{TP}{TP + FN} \tag{3}$$

High recall indicates that the model captures most of the relevant structure, reducing under-segmentation, which is important for completeness in tracing organelles.

### 2.4 Intersection over Union (IoU)

Also known as the Jaccard Index, IoU measures the fraction of intersection over union between prediction and ground truth:

$$\text{IoU}(P, G) = \frac{|P \cap G|}{|P \cup G|} = \frac{TP}{TP + FP + FN} \tag{4}$$

### 2.5 Variation of Information (VI)

VI is an information-theoretic metric that quantifies the distance between two clusterings (segmentations). Given a ground truth segmentation $G$ and predicted segmentation $P$, the VI is defined as:

$$\text{VI}(P, G) = H(P) + H(G) - 2I(P, G) \tag{5}$$

where $H(P)$ and $H(G)$ are the entropies of the predicted and ground truth segmentations, and $I(P, G)$ is the mutual information between them. Lower VI indicates higher similarity and better segmentation.
